# Condensation of Sidekick at tricellular junctions organizes mechanical forces for cell-cell adhesion remodeling

**DOI:** 10.1101/2025.04.25.650632

**Authors:** Hiroyuki Uechi, Daxiao Sun, Yuki Saeki, Tetsuya Hiraiwa, Alf Honigmann, Anthony A. Hyman, Erina Kuranaga

## Abstract

Epithelial morphogenesis relies on coordinated multicellular movements regulated by adhesion and force-generating molecules. It remains elusive how cells ensure organization of these molecules in space and time. Combining *in vitro* reconstitution and fly genetics, we demonstrate that condensation of an adhesive transmembrane protein shapes molecular dynamics at tricellular junctions to facilitate cell–cell junction remodeling. Specifically, we identify that the intracellular domain of Sidekick, known to localize to tricellular junctions and recruit mechanical force-generating myosin II, undergoes phase separation into condensates. This condensation controls the physiological localization and concentration of Sidekick on lipid membranes. Altering the molecular dynamics of these condensates disrupts Sidekick’s localization against cellular movements, misdirecting myosin II and leading to delayed junction formation. These findings suggest that cells exploit the mechano-resistant properties of tricellular junction-anchored condensates to spatially allocate force-regulating molecules. This study underscores the importance of condensation to organize tricellular junctions for robust multicellular movement.

## Introduction

Tissue morphogenesis is orchestrated by coordinated movements of the constituting cell collectives, such as cell division, deformation, migration, and rearrangement (intercalation)^1^. These multicellular behaviors are executed by proteins localizing at the plasma membrane such as cell adhesion and mechanical force-generating proteins^1–3^. Distribution and functioning of these molecules are tightly regulated at proper places; however, it remains elusive how dynamics of the morphogenesis-regulating molecules are ensured in space and time for multicellular coordination.

Biomolecular condensation regulates spatial compartmentalization and reaction dynamics of membrane-less organelles and molecular assemblies in the cytoplasm and the nucleoplasm^4^. Recent studies demonstrated that condensation also organizes dynamics of plasma membrane-localizing adhesive complexes such as the tight junction and the focal adhesion^5,6^, where proteins which form into condensates are mostly membrane-associated intracellular components like ZO-1 and Talin^7–10^. Condensates often generate their functions via their material properties, ranging from dynamic (liquid-like) to less dynamic (gel- or solid-like) states. Less dynamic condensates contribute to function against mechanical or external inputs, which was indicated on centrosomes^11^ and the Hippo signaling-mediating condensates^12^; in another case, less dynamic property engages in recruiting specific organelles and molecules to or through condensates, such as nuclear pores^13^, RNA-protein assemblies^14^, and Balbiani bodies^15^. Given the role in spatial organization, condensation could be physiologically relevant to dynamics of membrane-associating proteins driving collective cell movements.

To address this possibility, we focus on *Drosophila* germband extension, anteroposterior elongation of the embryonic ectoderm occurring just after onset of gastrulation^1,16–18^. This epithelial extension is induced by cell intercalation, in which cell–cell junctions along the dorsoventral axis shorten, and new junctions extend along the anteroposterior axis^17,19^. Such epithelial junction remodeling is organized at the adherens junction^20^, and associating actomyosin, the complex of actin and non-muscle myosin II, generates mechanical contractile forces to remodel junctions^20,21^ (introductory schema in Fig. S1A). Myosin II is also found in a meshwork structure, and myosin II meshwork in junction-forming cells flows from the medial-apical cortex to the adherens junctions to shorten the junctions while that in cells at the edges of new junctions flows to and “pulls” the vertices to extend the junctions (Fig. S1A)^19,22–24^. At the vertices (*i.e.*, tricellular junctions), the adhesive single-pass transmembrane protein Sidekick (Sdk) specifically localizes, is required for accumulation of myosin II, and then facilitates formation of new junctions^25–28^. However, it remains unexplored how Sdk enables itself to function at tricellular junctions to ensure junction remodeling.

In this study, we identify that the intracellular domain of Sdk forms into condensates, which can organize Sdk’s localization patterns and physiological concentration on the plasma membrane. We also show that the material property of the condensates underlies Sdk’s function: increasing the molecular dynamics of condensates by deleting an evolutionally conserved motif disturbs localization of Sdk and recruitment of myosin II, which delays junction extension. These results suggest a possible biophysical mechanism in which cells exploit the tricellular junction-anchored condensates to allocate mechanical force-regulating proteins, which eventually facilitates proper junction dynamics in collective cell movement.

## Results

### Condensation of the intracellular domain localizes Sdk to tricellular junctions

To probe which domain of Sdk is required to function at tricellular junctions, we used the CRISPR-Cas9 system to generate knock-in flies in which endogenous *sdk* is replaced with C-terminally mScarlet-tagged full-length wild-type Sdk (Sdk^WT^) or Sdk lacking its intracellular domain (Sdk^ΔICD^; Figs. 1A,B and S1B). Sdk^WT^::mScarlet and Sdk^ΔICD^::mScarlet showed comparable protein expression levels in fly pupae (Fig. S1C). They were imaged with endogenously GFP-tagged E-Cadherin (E-Cad::GFP)^29^ to label adherens junctions in the fly dorsal thorax epithelia at 24–26 h after puparium formation (APF), after the first round of cell division and at the late stage of cell deformation^30,31^, thus with less movement of junctions. This showed that, while Sdk^WT^::mScarlet was enriched at tricellular junctions similar to endogenous intact Sdk, Sdk^ΔICD^::mScarlet was observed in rather uniform distribution along junctions (Figs. 1B and S1D), suggesting that the intracellular domain is required to localize Sdk to tricellular junctions. This contrasts with another adhesive transmembrane protein Anakonda (Aka; Fig. 1A): this protein localizes to tricellular junctions at the level of septate junctions, which is preserved even when its intracellular domain is removed^32^.

**Figure 1.**
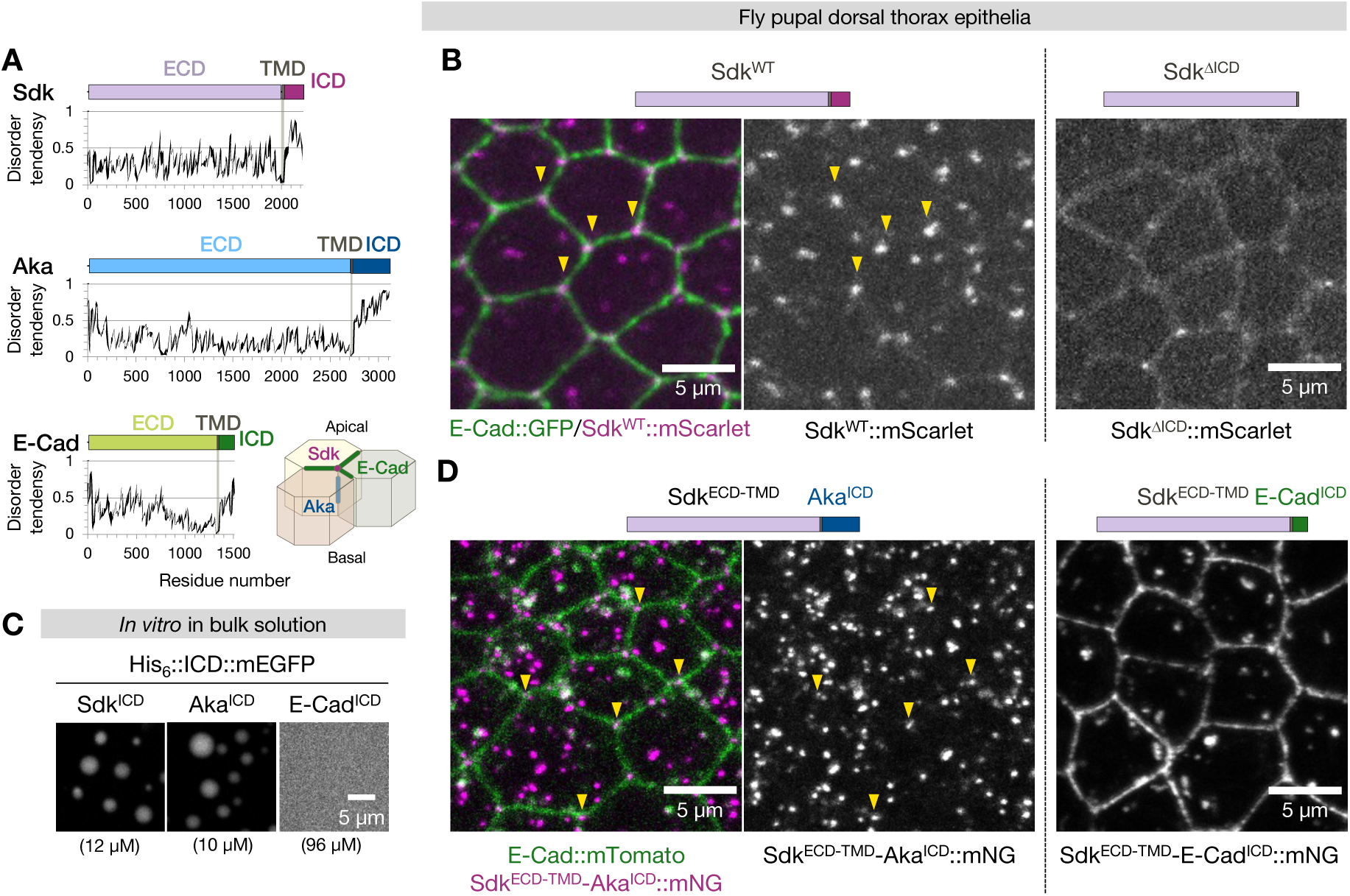
Condensation-mediated localization of Sdk at tricellular junctions. (A) Schema of the domain structures of *Drosophila* Sidekick (Sdk), Anakonda (Aka), and E-Cadherin (E-Cad), and their disorder prediction on IUPred3. ECD, extracellular domain; TMD, transmembrane domain; ICD, intracellular domain. Bottom right, schema of distribution of these adhesive transmembrane proteins in fly epithelial cells. Sdk and E-Cadherin are at the level of adheres junctions (sub-apical) while Aka is at the level of septate junctions (lateral). (B and D) Representative images of epithelial cells in the *Drosophila* pupal dorsal thorax endogenously expressing Sdk^WT^::mScarlet or Sdk^ΔICD^::mScarlet with E-Cad::GFP (to indicate bicellular junctions; (B)) or Sdk::mNG whose intracellular domain was replaced with that of Aka (Sdk^ECD-TMD^-Aka^ICD^::mNG) or E-Cadherin (Sdk^ECD-TMD^-E-Cad^ICD^::mNG) with E-Cad::mTomato (D) at 24–26 h after puparium formation (APF). Arrowheads, some of tricellular junctions. Note that the tagged proteins caused cytoplasmic aggregates as frequently observed with similarly tagged proteins expressed in fly pupal epithelia^25,63^. (C) Representative images of *in vitro* His_6_::ICD::mEGFP condensates in bulk phase (in buffer of 20 mM HEPES pH 7.25 and 150 mM KCl) at indicated bulk protein concentrations in the presence (Sdk^ICD^ and E-Cad^ICD^) and absence (Aka^ICD^) of 5% PEG-20,000. See also Figure S1.

Sdk intracellular domain (Sdk^ICD^) spans 200 residues and does not have sub-domains with known functions except a motif at the C-terminus called WAVE regulatory complex-interacting receptor sequence (WIRS) or the overlapping PDZ binding motif, which mediate protein–protein interactions^26,33^. Sdk is associated directly with Pyd (the orthologue of mammalian ZO-1) via the PDZ binding motif and indirectly with Cno (mammalian Afadin) possibly via yet unknown molecules^26^. Since ZO-1 and Afadin are demonstrated to form into condensates^7,34^, we first wondered if these proteins mediate the spotted localization of Sdk to tricellular junctions. However, although Pyd and Cno also have a preference to accumulate at tricellular junctions in epithelia as reported previously^26,33,35,36^(Fig. S1E), Sdk showed disproportionately higher enrichment to tricellular junctions in the dorsal thorax epithelia: the fold enrichment at tricellular junctions compared to at bicellular junctions was around 3.3 while that of Cno was 1.6 (Fig. S1D,E). Crucially, Sdk preserves its tricellular junction localization even when Pyd or Cno is depleted, although depletion of Cno could reduce Sdk levels concomitantly with reducing junctional E-Cadherin levels in some tissues^26^. Therefore, we speculated that Sdk’s localization is mediated by the intrinsic nature of Sdk^ICD^ or its not yet identified interacting molecules.

Instead of having structured sub-domains, Sdk^ICD^ largely comprises intrinsically disordered regions (IDRs; Fig. 1A, regions with score >0.5), which are known to be either driver or regulator of protein condensation^4,37^. To assess condensation behaviors of Sdk^ICD^, we purified a recombinant protein of N-terminally His_6_- and C-terminally monomeric enhanced green fluorescent protein (mEGFP)-tagged Sdk^ICD^ from insect cells. We observed that the purified protein (His_6_::Sdk^ICD^::mEGFP) formed into condensates in bulk phase (buffer solution in the test tube) at a saturation concentration of around 10 µM in the presence of a crowding agent polyethylene glycol (PEG; Figs. 1C and S1F).

To investigate if condensation is involved in Sdk’s localization in epithelia, we first utilized intracellular domains of other adhesive transmembrane proteins. A previous computational analysis of transmembrane proteins shows that IDRs are enriched at their cytoplasmic domains^38^. Indeed, the intracellular domain of Aka (Aka^ICD^) is disordered like Sdk, whereas that of E-Cadherin (E-Cad^ICD^) is less so compared to the others (Fig. 1A). We purified Aka^ICD^ and E-Cad^ICD^ in the same construct as Sdk^ICD^ and found that His_6_::Aka^ICD^::mEGFP formed into condensates in bulk phase at around 6 µM without PEG while His_6_::E-Cad^ICD^::mEGFP did not even at around 100 µM with PEG (Figs. 1C and S1F). We then endogenously replaced Sdk^ICD^ with these alternatives tagged with mNeonGreen (mNG) in flies and imaged the dorsal thorax epithelia (Figs. 1D and S1B). The chimeric protein with condensate-forming Aka^ICD^ (Sdk^ECD-TMD^-Aka^ICD^::mNG) showed spotted distributions on junctions, mostly found at tricellular junctions at the level of adherens junctions labeled with endogenously RFP-tagged E-Cadherin (E-Cad::mTomato) (Fig. 1D). This implies that the extracellular domain of Sdk contributes to confine the distribution to sub-apical regions. Meanwhile, the chimeric protein with diffusing E-Cad^ICD^ (Sdk^ECD-TMD^-E-Cad^ICD^::mNG) exhibited uniform distribution at junctions (Fig. 1D). To confirm that condensation itself is required for localization to tricellular junctions, we next swapped Sdk^ICD^ for another heterologous IDR not relevant to cell adhesion. The prion like domain (PLD) of human Fused in sarcoma (the N-terminal 211 amino acid residues; FUS^PLD^) is widely used for complementation experiments to test necessity of condensation by swapping IDRs of tested proteins for FUS^PLD^ (ref.^39^). The chimeric protein (Sdk^ECD-TMD^-FUS^PLD^::mScarlet) showed its signals at tricellular junctions (Fig. S1G). Collectively, these results suggest that Sdk localizes to tricellular junctions via condensation behavior of its intrinsically disordered intracellular domain.

### Sdk intracellular domain can recapitulate Sdk’s localization patterns and physiological concentration

To investigate direct contribution of Sdk^ICD^ condensation to Sdk’s distribution on the plasma membrane, we next analyzed intracellular domain proteins on *in vitro* lipid membranes. Protein condensation happens not only in 3D environments like the cytoplasm but also on surfaces such as the plasma membrane and DNA, where condensates can emerge around physiological protein concentrations often lower than the saturation concentration for bulk phase separation^40,41^. To test dynamics of Sdk^ICD^ on lipid membranes, we titrated His_6_::Sdk^ICD^::mEGFP proteins on supported lipid bilayers (SLBs) containing Ni-NTA-conjugated lipids to recruit the His-tagged proteins onto the SLBs, instead of introducing the transmembrane domain (Fig. 2A). On SLBs, His_6_::Sdk^ICD^::mEGFP started to form into small punctate structures at a bulk concentration of around 125 nM, two orders of magnitude lower than the bulk phase saturation concentration, suggesting lipid surface-assisted condensation of Sdk^ICD^ (Fig. 2B). The size of condensates on SLBs was smaller than that of previously reported focal adhesion and tight junction proteins^10,41,42^ but similar to the size of Sdk signals at tricellular junctions in epithelia (Figs. 1B and 2B). Such punctate structures were observed neither on SLBs devoid of Ni-NTA lipids nor with E-Cad^ICD^ (His_6_::E-Cad^ICD^::mEGFP), ruling out a possibility of non-specific binding of Sdk^ICD^ on lipid membranes (Figs. 2B and S2A). This minimal assay suggests that Sdk^ICD^ is sufficient to organize punctate distribution on lipid membranes.

**Figure 2.**
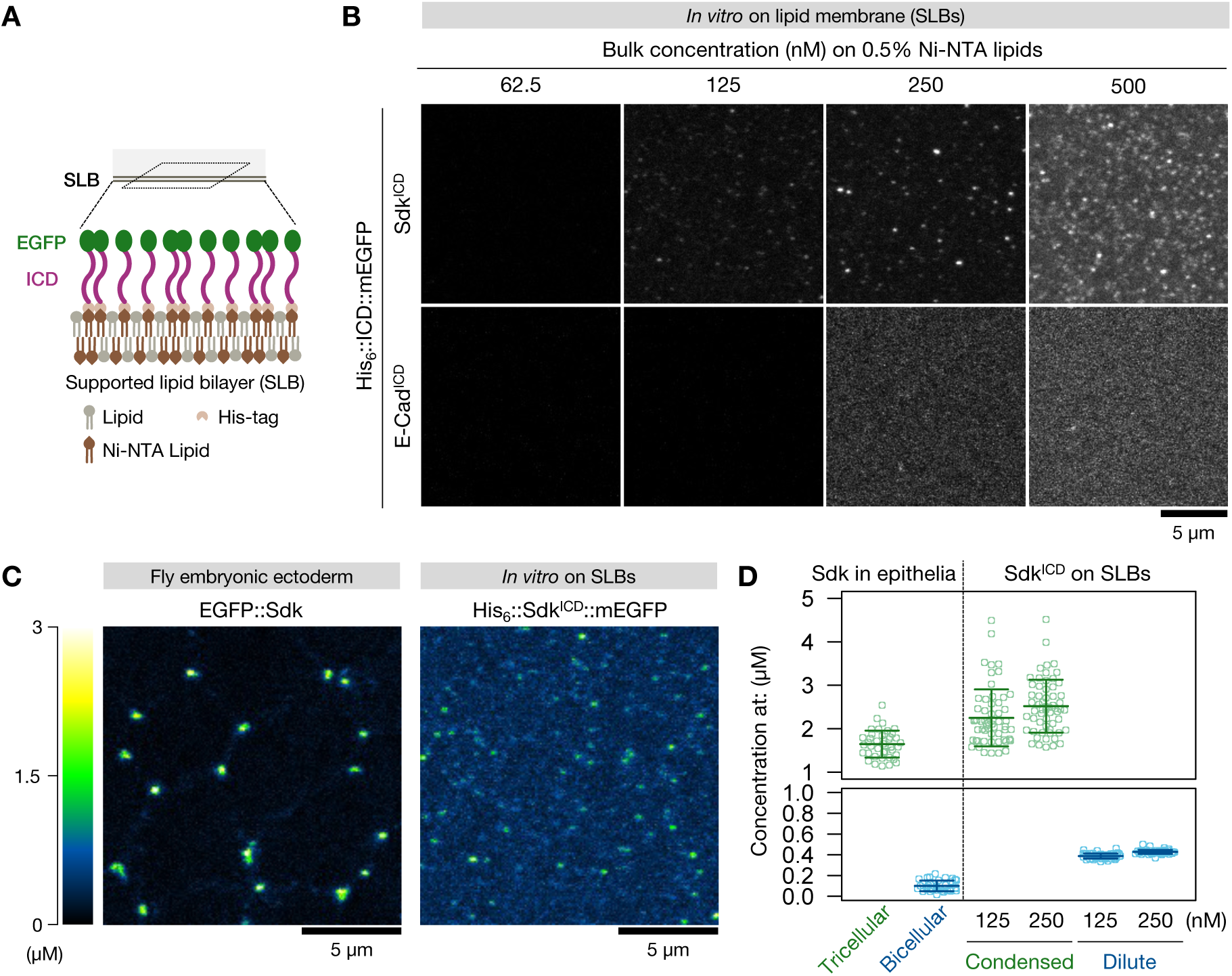
The intracellular domain of Sdk can shape Sdk’s localization and concentration. (A) Schema of *in vitro* protein condensation assay on supported lipid bilayers (SLBs). (B) Representative images of His_6_::Sdk^ICD^::mEGFP and His_6_::E-Cad^ICD^::mEGFP on SLBs containing 0.5% (molar ratio) Ni-NTA conjugated lipids in buffer (20 mM HEPES pH 7.25, 150 mM KCl, and 1 mM MgCl_2_) at indicated bulk concentrations. (C) Representative images of EGFP::Sdk in the ectodermal cells in fly embryos after germband extension (left) and His_6_::Sdk^ICD^::mEGFP at bulk concentration of 125 nM on 0.5% Ni-NTA lipid-containing SLBs (right; the same image as in (B)). The images are pseud-colored based on estimated protein concentrations. (D) Mean ± S.D. of the local concentrations of EGFP::Sdk in the ectoderm at tricellular or bicellular junctions and those of His_6_::Sdk^ICD^::mEGFP (125 and 250 nM bulk concentration) on SLBs (0.5% Ni-NTA lipids) at (condensed phase) and outside (dilute phase) the punctate structures. *n* = 41 tricellular and bicellular junctions each from four embryos for EGFP::Sdk; 60 regions of interest (ROIs) each phase for His_6_::Sdk^ICD^::mEGFP. See also Figure S2.

While the punctate Sdk^ICD^ distribution was observed uniformly on SLBs (Fig. 2B), *in vivo* Sdk shows dominant localization at tricellular junctions. We asked how the distribution is concentrated at such specific loci. The tricellular junction is composed of bent plasma membranes from three or more cells, with mostly convex angles. Since the intrinsically disordered state of Sdk^ICD^ suggests that this domain may exist in the shape of an unstructured polymer rather than a structured globule, such bent lipid membrane surface could make the associating Sdk^ICD^ closer to each other at the angled point, facilitate their homotypic interaction, and thus propel their condensation (Fig. S2B). To test if lipid geometry can affect dynamics of Sdk^ICD^, we introduced giant unilamellar vesicles (GUVs) on SLBs (Fig. S2C). After sedimentation, GUVs formed contact angles with SLBs at the contact edges (Fig. S2C,D), emulating lipid surface with convex angles. The proteins were then introduced onto this lipid mixture, which showed that His_6_::Sdk^ICD^::mEGFP but not His_6_::E-Cad^ICD^::mEGFP had higher signal intensity at the edges of lipid bilayer contacts compared to that on SLBs (Fig. S2D,E). This suggests that angled geometry of lipid membrane assists the localization of Sdk^ICD^, probably via facilitating condensation of Sdk^ICD^ at the angled position.

To further validate if the distribution of Sdk^ICD^ on SLBs is relevant to physiological Sdk distribution, we aimed to compare local concentrations of both Sdk^ICD^ on SLBs and Sdk in epithelia. To this end, we generated a standard curve by measuring signal intensities of purified mEGFP at known concentrations (Fig. S2F) and used a protein-trap line of *sdk* in which Sdk is endogenously tagged with the parental EGFP with an almost identical brightness to mEGFP (https://www.fpbase.org/) (EGFP::Sdk)^43^. Imaging of this embryonic ectoderm after germband extension with the same condition as the purified mEGFP inferred that the local concentration of EGFP::Sdk at tricellular junctions was 1.65 ± 0.31 µM (Fig. 2C,D). We then measured intensities of His_6_::Sdk^ICD^::mEGFP (bulk concentrations of 125 and 250 nM) on SLBs (0.5% Ni-NTA lipids), at which this protein started to form into punctate structures on the SLBs (Fig. 2B). The concentrations of His_6_::Sdk^ICD^::mEGFP at the punctate structures (condensed phase; 2.25 ± 0.66 and 2.52 ± 0.61 µM, respectively) were comparable to that of EGFP::Sdk at tricellular junctions (Fig. 2C,D). On the other hand, the concentrations of His_6_::Sdk^ICD^::mEGFP on SLBs outside the puncta (dilute phase) were 0.38 ± 0.024 and 0.43 ± 0.019 µM, respectively, smaller than that of EGFP::Sdk at tricellular junctions and rather close to that of EGFP::Sdk at bicellular junctions (0.10 ± 0.051 µM; Fig. 2C,D). This result suggests that condensation of Sdk^ICD^ on lipid membranes gives rise to Sdk’s concentration distribution in epithelia. Collectively, we propose that lipid membranes and condensation of the intracellular domain control localization of Sdk to tricellular junctions.

### A conserved amino acid motif encodes the material property of Sdk intracellular domain condensates

On SLBs, the size of His_6_::Sdk^ICD^::mEGFP condensates did not increase as a function of protein bulk concentrations (Fig. 2B), suggesting less dynamic property of this domain. To infer material properties of Sdk^ICD^ condensates, we used the condensates in bulk solution as they are larger than those on SLBs and thus allowed us to perform a fluorescence recovery after photobleaching (FRAP) assay on the half area of each condensate, which gives information of internal molecular rearrangement^44^. The half FRAP assay showed slow signal recovery at the bleached areas and almost unchanged signal intensity at the non-bleached areas, indicating slow molecular diffusion within the condensate (Fig. 3A. left). This result confirms that Sdk^ICD^ condensates are in less dynamic (*e.g.*, gel-like) state.

**Figure 3.**
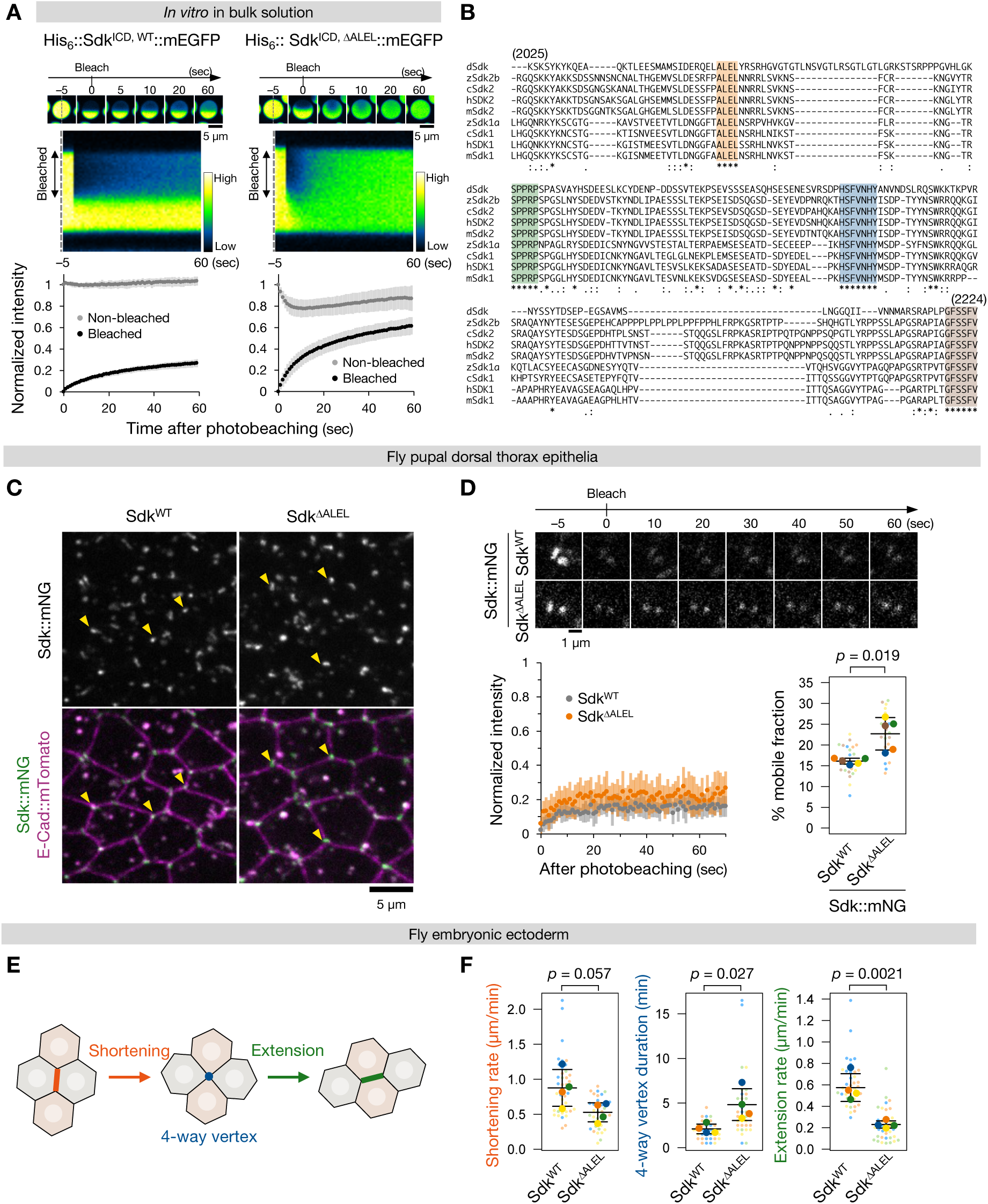
A motif encoding the material property of Sdk is involved in its molecular function. (A) Top, time-lapse images of one condensate each of His_6_::Sdk^ICD, WT^::mEGFP and His_6_::Sdk^ICD, ΔALEL^::mEGFP at 15 µM in bulk phase as in Figure 1C before and after photobleaching (0 sec) on the top half. The images are pseud-colored based on signal intensities. Middle, kymographs along broken lines on the condensates at the top (–5 sec). Bottom, mean ± S.D. of normalized intensities at bleached (black) and unbleached (grey) areas after photobleaching. *n* = 16 (Sdk^ICD, WT^) and *n* = 26 (Sdk^ICD, ΔALEL^) condensates. (B) Alignment of amino acid sequences of the intracellular domains of *Drosophila* Sdk and of Sdk1 and Sdk2 from vertebrates. Parentheses, positions of the first and last residues of *Drosophila* Sdk^ICD^. The very C-terminal motif contains the WIRS motif (FSSFV). c, chick; m, mouse; h, human; z. zebra fish. (C) Representative images of the pupal dorsal thorax epithelia endogenously expressing Sdk^WT^::mNG or Sdk^ΔALEL^::mNG with E-Cad::mTomato at 24–26 h APF. Note that images also show Sdk::mNG- and E-Cad::mTomato-positive cytoplasmic aggregates. (D) Top, representative images of Sdk^WT^::mNG or Sdk^ΔALEL^::mNG at two tricellular junctions each in the pupal dorsal thorax epithelia before and after photobleaching (0 sec). Bottom, mean ± S.D. of normalized intensity of each construct of Sdk::mNG after photobleaching (left) and of mobile fraction. *n* = 24 (Sdk^WT^) and 23 (Sdk^ΔALEL^) tricellular junctions from five flies each. *p*-value by an unpaired two-tailed Welch’s *t*-test. (E) Schema of junction remodeling in epithelial cell intercalation, composed of junction shortening (orange), formation and resolution of four-way vertices (blue), and extension of new junctions (green). (F) Mean ± S.D. of indicated junction dynamics during germband extension in Sdk^WT^::mScarelt or Sdk^ΔALEL^::mScarlet flies with E-Cad::GFP. *n* = 38 (Sdk^WT^) and 36 (Sdk^ΔALEL^) remodeling junctions from four flies each. *p*-values by an unpaired two-tailed *t*-test. See also Figure S3.

To examine the role of the less dynamic state, we screened mutants of Sdk^ICD^ with altered condensation behaviors. We reasoned that, if material property of Sdk^ICD^ is involved in Sdk’s function, such property may be encoded on evolutionally conserved amino acid residues. Amino acid sequences of IDRs are less conserved among orthologues in contrast to those of structured domains, especially when phylogenetically distant species like vertebrates and invertebrates are compared. Nevertheless, we found several motifs of four-to-seven successive amino acid residues which are completely identical between Sdk^ICD^ and the intracellular domains of its vertebrate orthologues Sdk1 and Sdk2, including the C-terminal WIRS motif/PDZ-binding motif (Fig. 3B). We purified recombinant proteins of Sdk^ICD^ devoid of each motif and generated their condensates, similarly to the wild-type intracellular domain (Fig. S1F). All the mutants formed into condensates in bulk phase at comparable saturation concentrations to the wild-type. However, FRAP on the whole condensate regions showed that condensates of one mutant lacking four amino acids, ^2054^Ala-Leu-Glu-Leu^2057^ (ALEL; His_6_::Sdk^ICD, ΔALEL^::mEGFP), had a higher mobile fraction compared to wild-type and the other mutant Sdk intracellular domains (Fig. S3A). The half FRAP assay on His_6_::Sdk^ICD, ΔALEL^::mEGFP condensates showed faster signal recovery at the bleached areas compared to His_6_::Sdk^ICD,WT^::mEGFP condensates, consistent to the whole FRAP assay (Fig. 3A). Signal intensity at the non-bleached area of His_6_::Sdk^ICD, ΔALEL^::mEGFP condensates decreased after bleaching, indicating molecular diffusion inside the condensates and thus dynamic (*e.g.*, viscoelastic) state. In agreement with the observed material differences, His_6_::Sdk^ICD, WT^::mEGFP condensates incubated over an hour did not relax after fusing while those of His_6_::Sdk^ICD, ΔALEL^::mEGFP did so (Videos S1,S2). These suggest that the less dynamic property of Sdk^ICD^ condensates is conferred by the ALEL motif.

On SLBs, condensates of His_6_::Sdk^ICD, ΔALEL^::mEGFP were formed at almost identical bulk protein concentrations and Ni-NTA lipid percentages to those of His_6_::Sdk^ICD, WT^::mEGFP but showed a slightly higher mobile fraction (Fig. S3B,C). Their mean mobile fractions were around 20–40%, analogous to those of other condensates on lipid membranes such as condensates of the focal adhesion protein Talin and the tight junction protein ZO-1^10,41^. His_6_::Sdk^ICD, ΔALEL^::mEGFP still accumulated at the edges of GUV-SLB contacts (Fig. S3D). Collectively, we identified a mutant of Sdk^ICD^ which has more dynamic molecular property than the wild-type but preserves condensation propensity and accumulation at angled positions of static lipid membranes.

### Flies of the mutant Sdk with increased dynamics phenocopy an *sdk* null mutant

We next turned back to fly epithelia to examine physiological roles of the dynamics of Sdk^ICD^. To this end, we generated knock-in flies of Sdk endogenously lacking the ALEL motif (Sdk^ΔALEL^) and tagged with the fluorescent proteins (Fig. S1B). Deleting the ALEL motif induced neither developmental delay nor lethality, in agreement with the previous observation that *sdk* null flies are viable^45,46^. In the dorsal thorax epithelia at 24–26 h APF, Sdk^ΔALEL^::mNG localized to tricellular junctions similar to Sdk^WT^::mNG (Figs. 3C and S3E). FRAP analysis of Sdk::mNG at tricellular junctions showed that a mobile fraction of Sdk^WT^::mNG was 16.11 ± 0.70% while that of Sdk^ΔALEL^::mNG was slightly higher (22.69 ± 3.91%; Fig. 3D). For both Sdk^WT^ and Sdk^ΔALEL^, the *in vivo* mobile fractions were smaller than the corresponding mobile fractions of His_6_::Sdk^ICD^::mEGFP on SLBs (Fig. S3C), probably reflecting inter-molecular interactions via Sdk extracellular domain at the plasma membrane. These observations indicate that the steady-state distribution and increased dynamics of this mutant characterized *in vitro* are preserved on the epithelial plasma membrane.

To validate effects of the ALEL deletion on multicellular dynamics, we examined junction remodeling during germband extension. Junction remodeling in epithelial cell intercalation is composed of shortening of junctions, formation of four-way vertices, and extension of new junctions after resolution of the four-way vertices (Fig. 3E)^1,17^. Depletion of *sdk* delays resolution of four-way vertices and extension of new junctions but shows almost no effect on junction shortening^25^. Analogous to this, comparison of germband extension between Sdk^WT^::mScarlet and Sdk^ΔALEL^::mScarlet flies showed that junction shortening rate was comparable between these strains, although Sdk^ΔALEL^ caused a marginal decrease with no statistical significance (Fig. 3F). In contrast, Sdk^ΔALEL^::mScarlet flies took longer time to resolve four-way vertices and showed reduced junction extension rate (Fig. 3F). This result suggests that the less dynamic property mediates Sdk’s ability to facilitate junction extension.

### The less dynamic state of Sdk confines its localization during cellular movement

To understand how mutating dynamics of Sdk^ICD^ condensates results in the defect of junction remodeling, we next examined molecular distribution of Sdk in germband extension. Epithelial cell rearrangement is accompanied by pulsatile contraction and expansion of junctions, which induces fluctuations of junction length and thus of the position of tricellular junctions^21,47^. Since distributions of Sdk^WT^ and Sdk^ΔALEL^ were comparable in the relatively static tissues (Fig. 3C), we wondered if this mutant shows altered distribution when junctions are dynamic. At tricellular junctions surrounded by non-remodeling junctions during germband extension, whereas both Sdk::mNG constructs mostly localized to the tricellular junctions, we observed dispersing or splitting of Sdk::mNG signals at the surrounding bicellular junctions (Fig. S4A). Most of the tricellular junctions did not show such dynamics in Sdk^WT^::mNG flies in an examined time window (10 min), while the abnormal dynamics were more frequent in Sdk^ΔALEL^::mNG flies with mostly one or two times of the dispersing event in 10 min (Fig. S4B). At remodeling junctions, Sdk is known to localize to not only tricellular junctions but also shortening and newly extending bicellular junctions at their late and early phases, respectively^25,27^. In addition to these dynamics, we observed dispersed dynamics of Sdk at the neighboring junctions of remodeling junctions, which emerged as discrete punctate or plaque-like signals of Sdk::mNG (Fig. 4A). During junction shortening, frequency of such dispersing events at neighboring junctions was mostly spanning from zero to two times in 10 min and was comparable between Sdk^WT^::mNG and Sdk^ΔALEL^::mNG flies (Fig. S4C). In contrast, the dispersing event was more frequent during junction extension in Sdk^ΔALEL^::mNG flies (spanning from one to six times in 10 min) compared to Sdk^WT^::mNG flies (mostly one or two times; Fig. 4B). This observation suggests that the less dynamic state of Sdk^ICD^ condensates is required to prevent abnormal localization of Sdk around tricellular junctions during multicellular movement and raised a possibility that altered Sdk dynamics underlies the defective junction extension in Sdk^ΔALEL^ flies.

**Figure 4.**
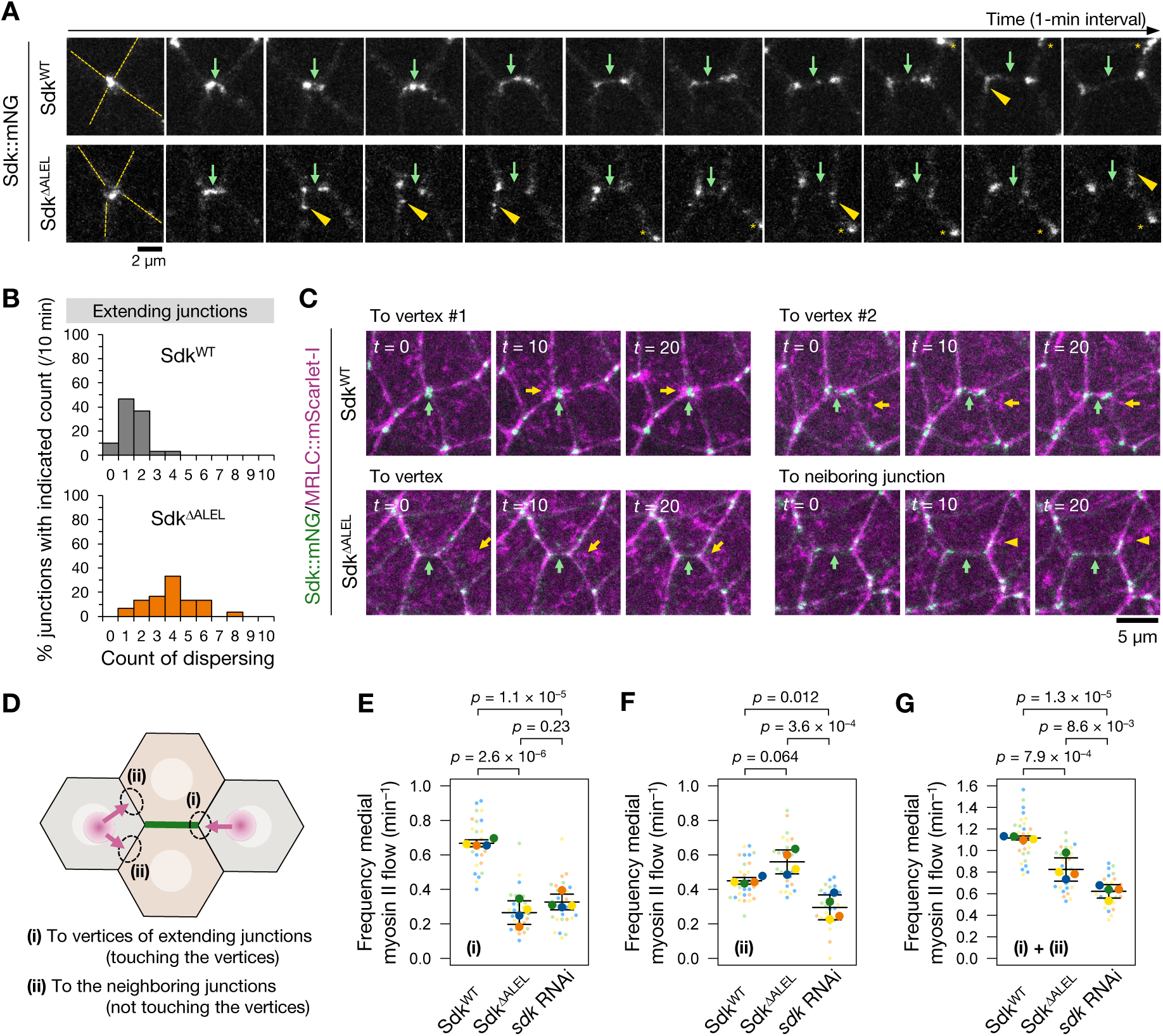
The less dynamic property of Sdk regulates molecular dynamics in junction remodeling. (A) Time-lapse images of Sdk::mNG around extending junctions from onset of four-way vertex resolution during germband extension. Broken lines, neighboring junctions (only labeled in the first images). Green arrows, the position of extending junctions. Yellow arrowheads, dispersed signals of Sdk::mNG on the neighboring junctions. Asterisks, Sdk::mNG at other tricellular junctions. (B) Proportion of frequency of the dispersing events of Sdk::mNG happening on the neighboring junctions of extending junctions within 10 min. *n* = 30 extending junctions from three embryos each genotype. *p* = 2.8 × 10^−8^ by two-tailed Mann-Whitney *U*-test. (C) Time-lapse images of Sdk::mNG and MRLC::mScarlet-I around extending junctions during germband extension. Two examples for each genotype. Green arrows, the position of extending junctions. Yellow arrows and arrowheads, medial-apical MRLC::mScarlet-I meshwork signals flowing to attach the vertices of extending junctions or the neighboring junctions, respectively. (D) Schema of medial apical myosin II flows (magenta) in the neighboring cells coming to (i) the vertices of extending junctions (green) or (ii) the neighboring junctions. (E to G) Mean ± S.D. of frequency of MRCL::Venus signals coming to the vertices (i; E), neighboring junctions (ii; F), and the sum ((i) + (ii); G) per min during cell– cell junction extension in germband extension. *n* = 29 extending junctions from four embryos each genotype. *p*-values by Tukey’s test. See also Figure S4.

### The mutant Sdk alters dynamics of the downstream molecule myosin II

To understand the link between the abnormal dynamics of Sdk and delay in junction extension, we first focused on mechanical force-generating myosin II. In the germ band, myosin II not only localizes to junctions but also appears as pulsative meshwork at cellular medial-apical cortices^19,22^. Such myosin II meshwork in neighboring cells at the edges of junctions flows to and “pulls” the vertices of the junctions, leading to junction extension (Fig. S1A)^19,24^. Since Sdk is required for myosin II accumulation at the tricellular junctions of extending junctions in another fly epithelial morphogenesis model^25^, we assumed that altered dynamics of Sdk may affect distribution of downstream myosin II during junction extension.

To test this possibility, we assessed myosin II dynamics by imaging myosin II regulatory light chain (MRLC) endogenously-tagged with mScarlet-I^48^ or Venus^49^ in the germ band with Sdk::mNG or Sdk::mScarlet, respectively (Fig. 4C–G). We also examined MRLC::Venus dynamics in *sdk*-depleted (*sdk* RNAi) flies to confirm that Sdk is required for recruitment of myosin II to tricellular junctions in germband extension (Fig. 4E–G). Consistent to the previous report, pulsative medial-apical MRLC::mScarlet-I meshwork signals in the neighboring cells of newly extending junctions frequently flowed towards the proximity of the vertices (tricellular junctions) of extending junctions (Fig. 4C, yellow arrows and arrowheads; Video S3)^19^. We noticed that the meshwork was eventually recruited to either the tricellular or neighboring junctions of newly extending junctions; the latter case was often concomitant with the dispersed localization of Sdk::mNG at the neighboring junctions (Fig. 4C, yellow arrowheads; Videos S3,S4). We categorized these MRLC meshwork dynamics into: (i) attachment to the tricellular junctions of newly extending junctions; (ii) no attachment to the vertices but instead to the neighboring junctions in the proximity (Fig. 4D). In *sdk*-depleted flies, frequencies of MRLC::Venus meshwork dynamics in both (i) and (ii) were decreased, supporting the idea that Sdk serves for myosin II recruitment to the tricellular junctions of extending junctions in germband extension (Fig. 4E,F). In Sdk^ΔALEL^::mScarlet flies, frequency of the MRLC::Venus meshwork recruited to the vertices was significantly decreased to a comparable level to *sdk* depletion (Fig. 4E). Meanwhile, Sdk^ΔALEL^::mScarlet flies showed marginal, although not statistically significant, increase in the frequency to be recruited to the neighboring junctions compared to Sdk^WT^::mScarlet flies, an opposite trend to *sdk*-depletion (Fig. 4F). This is possibly due to Sdk^ΔALEL^ mis-localizing to the neighboring junctions. Overall, frequency to be eventually attached to around the tricellular junctions (sum of (i) and (ii)) was decreased in Sdk^ΔALEL^::mScarlet flies, analogous to *sdk*-depleted flies (Fig. 4G).

We next aimed to examine Pyd and Cno as the other known Sdk-associating molecules^26^. However, Pyd::EGFP signals (endogenously EGFP-tagged Pyd in its protein trap line) during germband extension were too weak to validate its spatiotemporal distribution in this tissue movement. Meanwhile, Cno is detectable during germband extension^35,50^, and its levels at tricellular junctions influence the four-way vertex resolution in cell rearrangement^35^. Using endogenously Venus-tagged Cno, we observed that once decreased Cno::Venus levels at four-way vertices gradually increased at the tricellular junctions of extending junctions in Sdk^WT^::mScarlet flies, consistent to a previous observation (Fig. S4D)^35^. In Sdk^ΔALEL^::mScarlet flies, such re-accumulation of Cno::Venus during junction extension tended to be attenuated (Fig. S3D). This suggests that altered Cno dynamics may also underlie the defective junction extension in Sdk^ΔALEL^ flies. We have to note that, since tagging close to a PDZ-binding motif can affect association with its binding proteins^51^, we might underestimate dynamics of Cno::Venus in the Sdk::mScarlet flies by altering dynamics of a possible PDZ-containing protein which links Cno to Sdk^26^. Further detailed studies are needed to probe cooperation of Cno on Sdk-mediated junction extension.

Collectively, these results suggest that Sdk recruits myosin II meshwork to tricellular junctions via its less dynamic condensates, which is required for efficient junction remodeling.

## Discussion

Biomolecular condensation underlies mechanisms for intracellular compartmentalization and dynamics of biomolecules. This biophysical property has also been demonstrated to organize protein assemblies on the cytoplasmic side of the plasma membrane such as adhesion complexes and signaling complexes^5^, for which most studies focus on condensation of cytoplasmic proteins. Meanwhile, the intracellular domain of the adhesive transmembrane protein Nephrin is shown to be capable of forming condensates in the presence of oppositely charged partner proteins in the test tube^52^, implying that condensation can also underlie dynamics of transmembrane proteins. Our study extends this concept by demonstrating that Sdk undergoes condensation via its intracellular domain autonomously, rather than through cytoplasmic partner proteins. Our results indicate that lipid membrane-assisted condensation of Sdk intracellular domain is necessary and sufficient for establishing the molecular localization patterns and concentration of Sdk in epithelia.

Our analyses of the ΔALEL mutant show that increasing dynamics of Sdk intracellular domain condensates disturbs to maintain localization of Sdk to tricellular junctions when junctions are dynamic, such as fluctuating in length and undergoing remodeling. Since these junctional dynamics are induced by contractile forces of actomyosin, this observation suggests that the material property of condensates ensures Sdk’s specific localization against mechanical forces. Such mechano-resistance of condensates deployed by less dynamic or gel/solid-like states is proposed or demonstrated on other subcellular compartments such as the centrosome (against pulling forces of microtubules)^11^, the focal adhesion (against contractile forces of actomyosin)^10^, and Merlin condensates in the Hippo signaling (against cytoskeletal tension)^12^. The following results suggest that maintaining Sdk to tricellular junctions consequently directs dynamics of downstream myosin II, allowing myosin II to flow to the tricellular junctions of extending junctions and thus leading to robust junction extension. Given that less dynamic condensates have been implicated in stabilizing protein localization under mechanical stresses, our results reinforce the idea that the mechano-resistance by Sdk condensation serves for proper distribution of mechanical forces in epithelial cell rearrangement. However, we have to note that Sdk could indirectly modulate myosin II dynamics via other downstream molecules. Collectively, these findings suggest that Sdk exploits its condensation with less dynamic state to withstand while allocate mechanical forces at tricellular junctions for collective cell movement. More broadly, our study proposes that adhesion proteins can self-organize into condensates at interfaces between multiple cells, influencing junctional mechanics.

The ALEL motif is conserved across species, which implies that the material property might be also evolutionally conserved and utilized to function. However, it is possible that the ALEL motif serves as a possible protein binding site, similar to the C-terminal WIRS/PDZ-binding motif. It remains unclear how this motif generates the observed physical chemical property. Alanine is shown to decrease a mobile fraction of the protein condensates when its repeat is expanded^53^; a familial amyotrophic lateral sclerosis-linked mutant FUS in which one glycine is substituted to glutamate is revealed to form into less dynamic condensates than wild-type FUS condensates^54^. It is possible that similar physical chemical properties underlie the ALEL motif-mediated material state of Sdk.

Our small survey of condensation propensity showed that Sdk is not an only adhesive transmembrane protein whose intracellular domain can autonomously undergo condensation: another protein Aka also had the condensate-forming intracellular domain with IDRs. We noticed some other intracellular domains with IDRs of fly adhesive single-pass transmembrane proteins form into condensates *in vitro* (bulk phase) whereas those with less IDRs did not (Fig. S4D), analogous to the observation on Sdk, Aka, and E-Cadherin intracellular domains. This notice could lead to an idea that other adhesive transmembrane proteins with IDR-enriched intracellular domains may also generate their dynamics and functions for cellular adhesion, mechanotransduction, and movement in part via condensation. Also, the differences of condensation propensity could underlie the reason why cells deploy a variety of adhesive transmembrane proteins. For example, since intracellular domains of tricellular junction proteins tested (Sdk, Aka, and Gliotactin [Gli]^55^) all formed into condensates *in vitro* (Figs. 1C and S4D), it is possible that cells chose to have the condensation-prone transmembrane proteins in order to give the identity to tricellular junctions, while proteins with less condensation propensity like cadherin proteins were chosen to meet the need for broadly ensuring adhesion along cell–cell interfaces.

## Resource availability

### Lead contact

Requests for further information and resources should be directed to and will be fulfilled by the lead contact, Hiroyuki Uechi.

### Material availability

All unique materials generated in this study are available from the lead contact upon request.

### Data and code availability

All the data reported in this study will be shared by the lead contact upon request. This study does not report original code.

## Methods

### Fly genetics

The following *Drosophila* strains were used except for those newly generated in this study: *E-Cad::GFP* (RRID: BDSC_60584)^29^; *E-Cad::mTomato* (RRID: BDSC_58789)^29^; *EGFP::Sdk* (RRID: BDSC_60169)^43^ *MRLC::mScarlet-I* (RRID: BDSC_94929)^48^; *MRLC::Venus* (FBal0360412)^49^; *Cno::Venus* (RRID: DGGR_115111); *Pyd::EGFP* (RRID: BDSC_60166); *mat*α*Tub-Gal4VP16^67C^* (RRID: BDSC_80361); *UAS-sdk RNAi^HMS^*^00292^ (RRID: BDSC_33412); *vas-Cas9* (RRID: BDSC_51324); *Crew* (RRID: BDSC_1092).

Flies were raised, crossed, and imaged at 25°C. Detailed genotypes in each experiment are listed below.

### Generation of knock-in flies

The series of Sdk fly strains were generated by CRISPR-Cas9 system-mediated homologous recombination^56^ as described with minor modifications^25,49^. In brief, for C-terminally tagged Sdk^WT^ knock-in flies, the oligonucleotide (5-G-TCAGACGAATGACGAGAAGC-3), corresponding to the sequence just before 3’ UTR in the last exon of *sdk*, was cloned into the pBFv-U6.2 vector (AddGene #138400) as the guide RNA (gRNA) plasmid (hereafter referred to as gRNA^WT^ plasmid). For the targeting vectors, the Venus cDNA sequence in the pPVxRF3 vector (AddGene #138385) was firstly replaced with cDNAs of either mNeonGreen or mScarlet (the sequences were codon-optimized in Max Planck Institute of Molecular Cell Biology and Genetics, Dresden, Germany) while the RFP (DsRed-Express2) sequence under the eye-specific promotor 3xP3^57^, used as a selection marker, and the flanking loxP sites were kept. These targeting vectors are referred to as pPmNGxRF3 and pPmSxRF3, respectively. The *sdk* genomic sequences before (500 bp; 5’ arm) and after (500 bp; 3’ arm) the stop codon, the former of which contained a silent mutation on the NGG sequence, were then cloned into pPmNGxRF3 and pPmSxRF3 vectors with KpnI and AscI sites, and SacII sites, respectively, with no frame shift between the last exon and the cDNAs of mNeonGreen/mScarlet.

For C-terminally tagged Sdk^ΔICD^ knock-in flies, the oligonucleotide (5-G-AAAGACTCTAGAGGAATCTA-3), corresponding to the sequence encoding the beginning of Sdk intracellular domain, was cloned into the pBFv-U6.2 vector as gRNA^ΔICD^ plasmid and used in combination with gRNA^WT^ plasmid. For the targeting vector, the 5’ arm of the Sdk^WT^ targeting vector was replaced with 486 bp of the *sdk* genomic sequences which contain the sequence encoding the first 14 amino acid residues of Sdk intracellular domain (2025–2038 a.a.) and a part of the gRNA^ΔICD^ sequence.

For ICD-swapped knock-in flies, the same gRNA plasmids for Sdk^ΔICD^ were used. For the targeting vectors, the 5’ arm of the Sdk^ΔICD^ targeting vector was flanked by cDNA sequences encoding intracellular domains of either Aka or E-Cadherin without each stop codon or the prion-like domain of human FUS.

For C-terminally tagged Sdk^ΔALEL^ knock-in flies, the oligonucleotide (5-G-CGAGCGACAAGAGTTGGCTC-3), corresponding to the *sdk* genomic sequence which contains a part of the sequence encoding the ALEL motif, was cloned into the pBFv-U6.2 vector as gRNA^ΔALEL^ plasmid and used in combination with gRNA^WT^ plasmid. For the targeting vectors, the 5’ arm of the Sdk^WT^ targeting vectors was replaced with 1182 bp of the *sdk* genomic sequences before the stop codon which contains the silent mutation on the NGG sequence for gRNA^WT^ and excludes the 12-bp sequence encoding the ALEL motif.

All the targeting vectors and gRNA plasmids were injected into *vas-Cas9* flies, which was performed by BestGene Inc. The selection marker DsRed-Express2 was removed when needed by crossing with Cre recombinase-expressing flies.

### Expression and purification of recombinant proteins

cDNAs of intracellular domains were cloned into the pOCC vectors^58^ such that the ICDs were tagged with 6×His (His_6_), monomeric EGFP, and maltose binding protein (MBP). The His tag was put at the transmembrane domain side while mEGFP and MBP were at the opposite terminus. Deletion of amino acids was performed with the standard mutagenesis procedure. Baculoviruses that can express the recombinant proteins were generated in Sf9 cells in a way established previously^58^. Protein purification was performed as described previously^59,60^. In brief, the expressed recombinant proteins (His_6_::ICD::mEGFP::MBP or MBP::mEGFP::ICD::His_6_) were extracted from either Sf9 or T.ni cells, in a buffer containing 20 mM HEPES pH 7.25, 1 M KCl, 5% glycerol, 1 mM DTT, and 20 mM imidazole supplemented with protein inhibitor cocktail (cOmplete, EDTA-free; Roche), and subjected to affinity purification using HisTrap HP (Cytiva) and amylose resin (New England Biolabs). The MBP tag was cleaved out by 3C protease, and the target proteins were concentrated by dialysis and further purified by size exclusion chromatography. The proteins were snap-frozen and stored in the storage buffer (20 mM HEPES pH 7.25, 500 mM KCl, 5% glycerol, and 1 mM DTT).

### Microscopic settings for imaging

Imaging was performed at 25°C throughout the manuscript. *In vitro* protein dynamics were imaged on either a TiE inverted microscope (Nikon) with an Apo 60× NA 1.2 water immersion objective (Nikon), a CSU-X1 spinning disk (Yokogawa), and an iXon EM+ DU-897 EMCCD camera (Andor) or LSM980 (Zeiss) with a 63× Plan-Apochromat NA 1.4 oil immersion objective (Zeiss). xy images of proteins in bulk solution are from single z section. xy images of proteins on SLBs are maximum-intensity projections of two to three serial optical sections taken at a 0.5-µm z step size. Time-lapse images were acquired at a 1-sec interval.

*In vivo* (fly) protein and cellular dynamics were imaged on either a IX83 ZDC inverted microscope (OLYMPUS) with a 63× UPlan SApo NA 1.3 silicone immersion objective (OLYMPUS), a CSU-W1 spinning disk (Yokogawa), and an iXon Ultra 888 monochrome EMCCD camera (Andor) or LSM980 as described above. All time-lapse images are maximum-intensity projections or single sections at the level of the adherens junctions taken at a 0.5-µm z step size. Time-lapse images were acquired at a 30-sec interval for E-Cadherin (cell–cell junction) and Sdk dynamics; a 10-sec interval for MRLC and Cno dynamics.

### Sample preparation for imaging flies

For imaging the fly dorsal thorax epithelia at the pupal stage, pupae 24 to 26 h APF were dissected and mounted as described previously^61^ except for covering the pupae: pupae were surrounded with silicon and covered with water and a cover glass.

For imaging germband extension, fly embryos before stage 7 were dechorionated with 5% sodium hypochlorite for 30 sec and washed with water, and then the ventrolateral side was attached onto the bottom of glass-bottom dishes.

### *In vitro* condensation assay in bulk solution

Proteins in storage buffer were diluted with HEPES pH 7.25 and PEG-20,000 to give 20 µl of indicated concentrations of the proteins, 20 mM HEPES pH 7.25, and 150 mM KCl in the presence or absence of 5% (w/v) PEG-20,000 and placed on a 384-well plate (Greiner #781096). The plate was Incubated at 25°C for 30 min to 1 h before imaging.

### *In vitro* condensation assay on lipid membrane

For protein dynamics on supported lipid bilayers (SLBs), the planar lipid membrane was prepared as described previously^41,62^ with minor modifications. Briefly, phospholipids composed of 1-palmitoyl-2-oleoyl-glycero-3-phosphocholine (16:0-18:1 PC or POPC; Avanti #850457), 0.25, 0.5, 0.75, or 1% (molar ratio) 1,2-dioleoyl-sn-glycero-3-[(N-(5-amino-1-carboxypentyl)iminodiacetic acid)succinyl] (nickel salt) (18:1 DGS-NTA(Ni); Avanti #790404), 0.1% 1,2-dioleoyl-sn-glycero-3-phosphoethanolamine-N-[methoxy(polyethylene glycol)-5000] (ammonium salt) (18:1 PEG5000 PE; Avanti #880230), and 0.1% 1,2-dioleoyl-sn-glycero-3-phosphoethanolamine-N-(Cyanine 5) (18:1 Cy5 PE; Avanti # 810335) in 100 µl of chloroform were put in a glass vial and dried up under vacuum at room temperature for 2 h. The dried lipids were suspended in 500 µl of buffer (20 mM HEPES pH 7.25, 150 mM KCl, and 1 mM MgCl_2_) to set 1 mM of the lipids and shaken at 200 rpm at 37°C for 2 h. The lipid mixture was transferred to a 1.5 ml tube and subjected to 15 times of freeze (in liquid nitrogen)-thaw (in 42°C water bath) cycles to generate small unilamellar vesicles (SUVs). The SUV mixture was clarified by centrifugation at 17,000 × *g* for 30 min, and the supernatant containing the SUVs was stored at 4°C. To make SLBs, a glass-bottom 384-well plate (Greiner #781892) was firstly cleaned with 2% Hellmanex II (Hellma) overnight and then 6 N NaOH for 30 min twice at room temperature, and the wells were washed with water and the buffer several times each. 10 µl of the SUV mixture was added to 30 µl of the buffer in each well and incubated at 25°C for 1 h, followed by addition of 10 µl of 5 M NaCl at 25°C for additional 1 h to collapse the SUVs onto the glass bottom. The wells were washed eight times with 50 µl of the buffer to remove unbound lipids with always leaving 50 µl of buffer on SLBs. The SLBs were blocked with 1 mg/ml BSA in the buffer at 25°C for 30 min. Then proteins were added (less than 1% volume) onto the 50 µl of the SLB containing buffer to set the final bulk concentrations and incubated at 25°C for 1 h before imaging.

For protein dynamics on SLBs and giant unilamellar vesicles (GUVs), 4 mM of phospholipids with the same composition as SLBs in chloroform was put on a 4% polyvinyl alcohol (PVA)-coated glass slide chamber and dried up under vacuum at room temperature for 1 h. Then 400 µl of the buffer was added to dried lipids, and the chamber was incubated at 40°C for 1 h to cause swelling of lipids, thus generating GUV-containing buffer. The GUV-containing buffer was added to a glass-bottom 384-well plate containing SLBs and the buffer to get the total buffer volume of 50 µl/well and incubated at 25°C for 1 h to settle down the GUVs onto the SLBs. The lipid mixture was blocked, mixed with proteins, and imaged as described above. Signal intensities were analyzed on the Fiji software (https://fiji.sc/) with manually set region of interests (ROIs).

### Measurement of local protein concentration in fly tissues and on lipid membrane

To generate a standard curve, purified mEGFP proteins at known concentrations in a range from 0 to 2.56 µM were imaged without signal saturation. The obtained mean pixel intensities were plotted as a function of protein concentrations and linear-fitted on Excel as a standard curve. EGFP-tagged proteins in the fly ectoderm after germband extension and on SLBs were imaged on the same microscope and same setting as those for mEGFP proteins. These intensities were measured on the Fiji software with manually set ROIs. The corresponding concentrations were estimated with the mean intensity inside the ROIs and the standard curve.

### Fluorescent recovery after photobleaching (FRAP)

FRAP was performed on either the CSU-X1 or the LSM980 systems. Time-lapse images of signals of mEGFP- or mNeonGreen-tagged proteins were acquired at a 1-sec interval both before (5 frames) and after (60 to 115 frames) photobleaching with a 488 nm laser. ROIs of a photobleached and a reference region were manually set based on outlines of each signal (that on condensates or at tricellular junctions). The images were analyzed on the Fiji software using a Python script available online (https://imagej.net/tutorials/analyze-frap-movies).

### Immunoblotting

Fly pupae were homogenized in buffer containing 20 mM HEPES (pH 7.25), 0.5% Triton X-100, and 150 mM NaCl and clarified by centrifugation at 20,000 × *g* for 10 min at 4°C. The samples were separated by SDS-PAGE, transferred to a PVDF membrane, and subjected to an immunoblot analysis with the following antibodies for immunoblotting (all were diluted at 1:5000): mouse anti-RFP (6G6; Chromotek) and mouse anti-α-Tubulin (DM1A; Sigma).

### Quantification of remodeling junction dynamics

Junction dynamics were analyzed manually using the Fiji software as done previously with minor modifications^25^. In brief, the shortening and growth rates were estimated with a linear fit of the length of junctions labeled with E-Cad::GFP measured every 30 sec for a total of 5 min before and after reaching 0.6 µm, respectively. The 4-way vertex duration time was defined when remodeling junctions were below 0.6 µm in length, and the duration of the 4-way vertex phase was measured from time-lapse images captured at 30-sec intervals.

### Quantification of *in vivo* molecular dynamics

Molecular dynamics were tracked manually on the Fiji software. For Sdk dynamics, emergence of ectopic Sdk::mNeonGreen signals (puncta or plaque-like structures) on bicellular junctions was counted in 10 min, and the proportion of frequency was calculated.

For myosin II dynamics, flows of MRLC meshwork signals at medial-apical cortices imaged for 4–22 min were categorized based on the direction: (i) those which attached to the vertices of extending junctions; (ii) those which did not attach to the vertices but instead to the proximity on the neighboring junctions. Those attaching to elsewhere on junctions or not attaching to junctions were not counted.

For Cno dynamics, Cno::Venus signals at tricellular and bicellular (the neighboring) junctions were measured with manually set ROIs (circles and lines, respectively).

### Statistics

Statistical analyses were performed on the R software or Excel. The statistical tests used for each analysis as well as the sample sizes are indicated in the corresponding figure legend. For those which have experimental replicates, tests were performed using each mean (bold bigger dots) from individual scores (pale smaller dots) within the corresponding replicate.

### List of imaging conditions

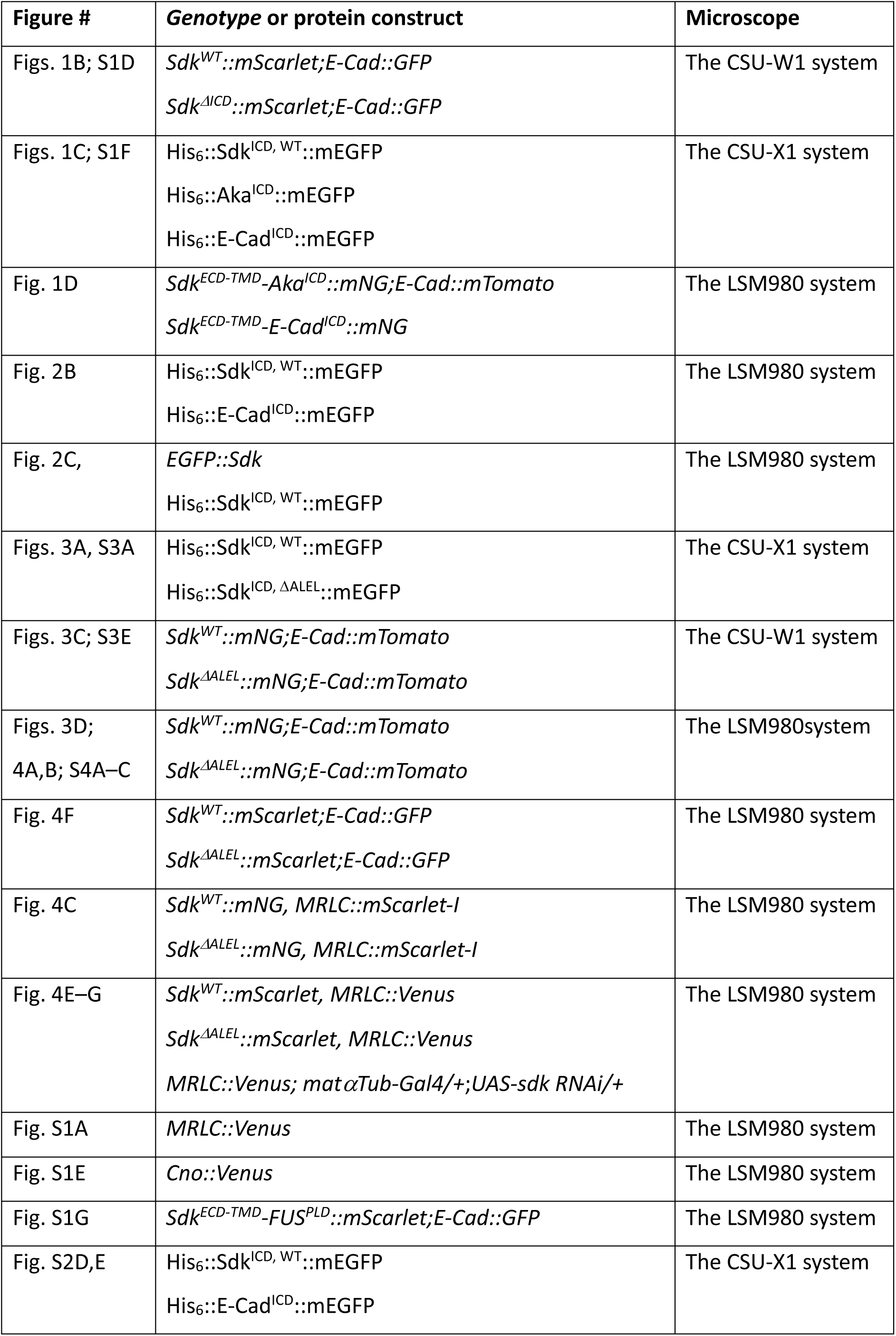
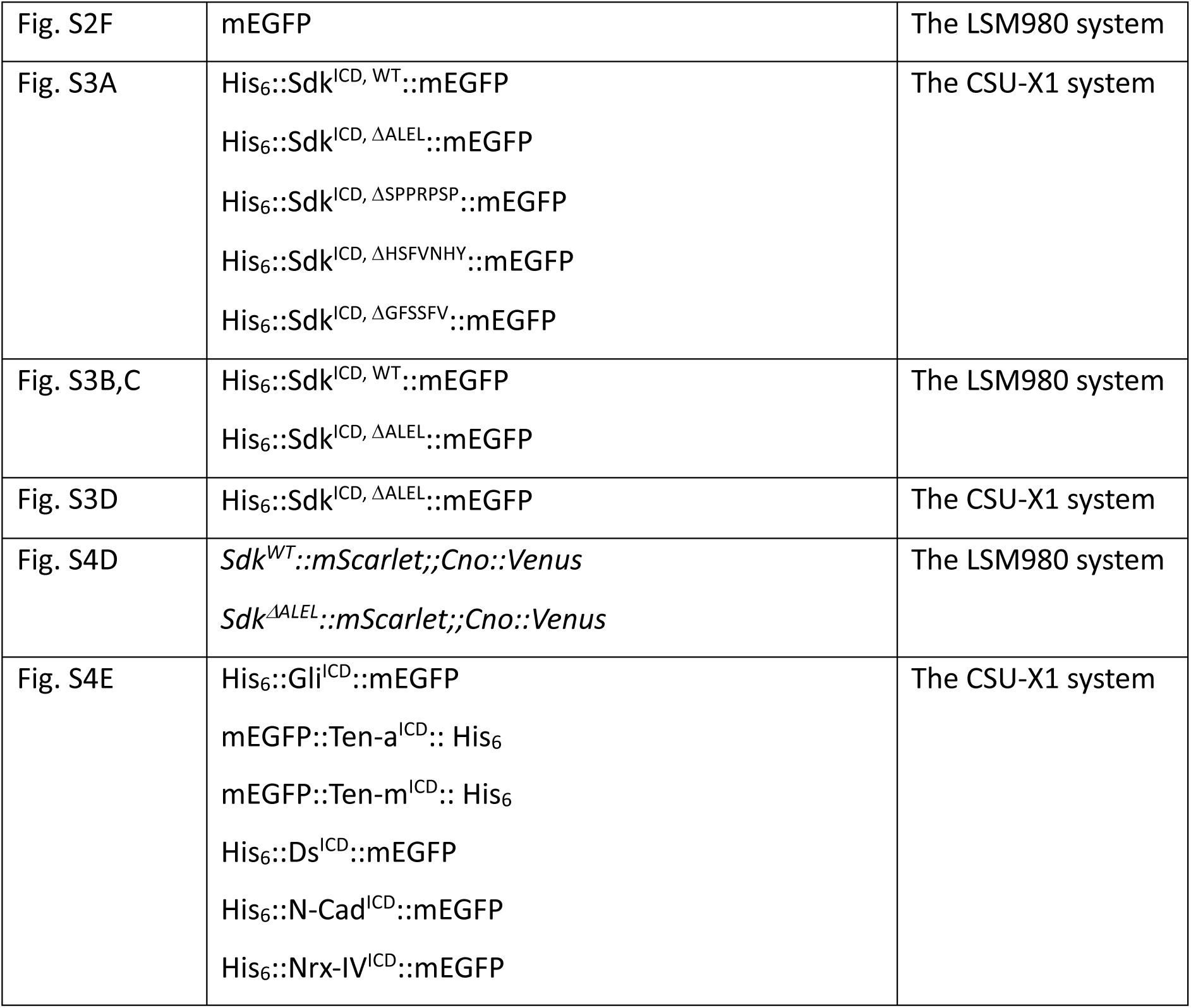

## Acknowledgements

We thank Bloomington Drosophila Stock Center and Kyoto Drosophila Stock Center for the fly stocks; Ilka Reichardt-Gómez for sharing cDNA sequences of codon-optimized mNeonGreen and mScarlet; Daiki Umetsu for providing the fly strain of *MRLC::Venus*; Carsten Höge for valuable review comments on the manuscript; the following facilities for their support: fly team, LMF, and PEPC (MPI-CBG), and the FRIS CoRE (Tohoku University). This study was supported by grants from MEXT/JSPS Grants-in-Aid for Scientific Research (KAKENHI) grant numbers JP23K19361 (H.U.), JP24K09448 (H.U.), JP24H01344 (H.U.), JP21H05255 (E.K.), and 24H00564 (E.K.); JST CREST grant number JPMJCR1852 (E.K.); AMED moonshot grant number 22zf0127001h002 (E.K.); The Uehara Memorial Foundation (H.U.).

## Author contribution

H.U. conceived the study. A.A.H. provided directions to the study. H.U. performed all the experiments with help from D.S. (*in vitro* assays with lipid membranes) and Y.S. (live imaging of fly embryos). D.S., T.H., and A.H. provided critical insights for interpreting the results. H.U. wrote the manuscript with review from D.S., T.H., A.H., A.A.H., and E.K. H.U. and E.K. acquired the fundings.

## Supplemental information

**Figure S1.**
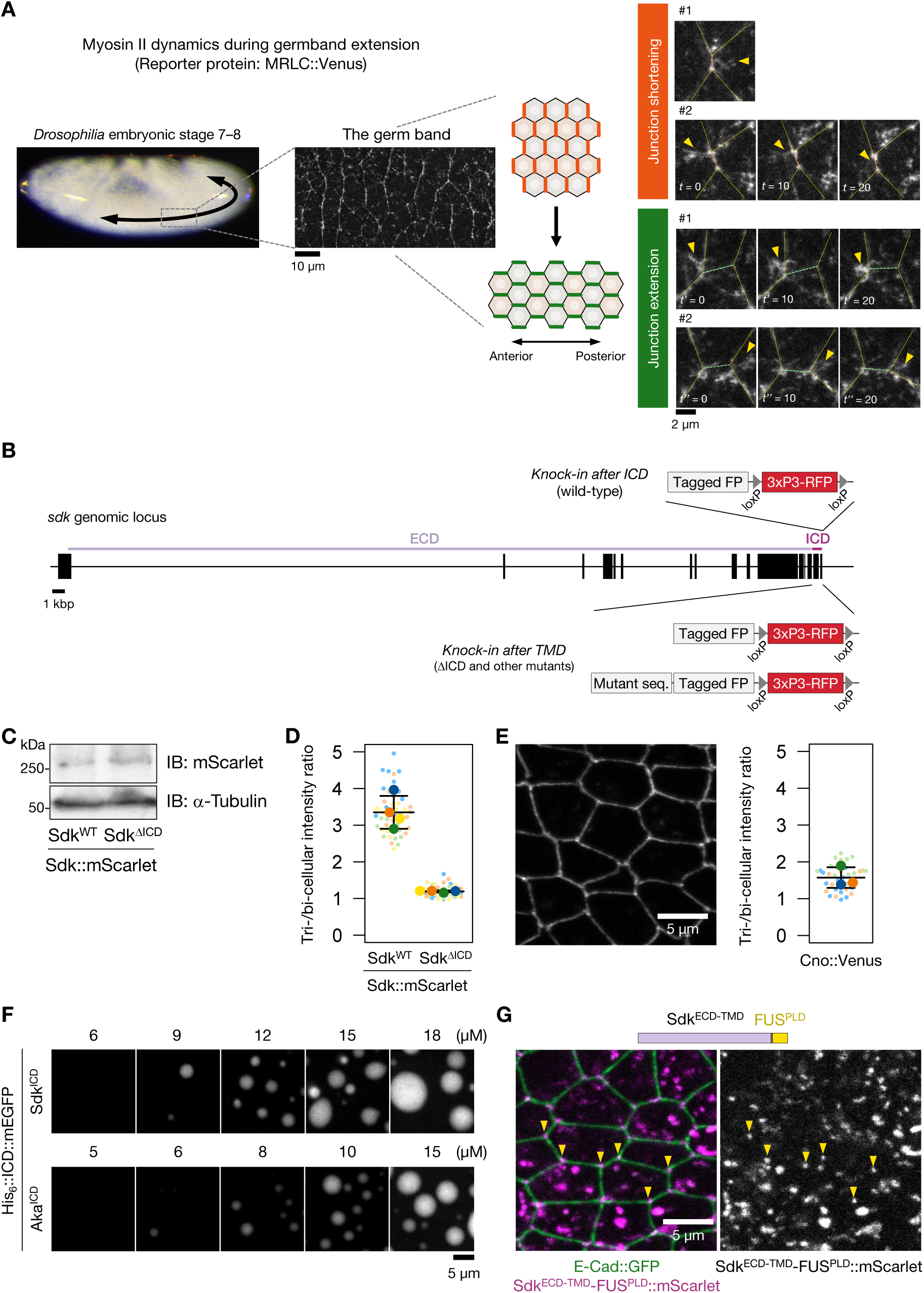
*In vivo* and *in vitro* molecular dynamics related to Figure 1. (A) Schematic introduction of myosin II dynamics underlying body axis elongation during *Drosophila* early embryogenesis: germband extension (references in the main text). This is anteroposterior extension of the ectoderm happening concomitantly with gastrulation around embryonic stage 7 (left). At the tissue level, the ectoderm shows polarized distribution of myosin II (indicated with Venus-tagged myosin II regulatory light chain [MRLC::Venus]) at junctions to induce shortening of junctions vertical to the anterior-posterior axis (middle). At the cellular level, in addition to junctions, myosin II distributed to medial apical cortices in a meshwork structure (right, arrowheads) which flows to shortening junctions (orange) and the vertices of extending junctions (green), driving junction remodeling-mediated epithelial cell intercalation. (B) Schema of the genomic locus of *sdk*. For knock-in flies, the tag and selection marker (see Methods) were inserted around the Sdk intracellular domain-encoding region. (C) Immunoblotting of pupal extracts from Sdk^WT^::mScarlet or Sdk^ΔICD^::mScarlet flies with indicated antibodies. (D) Mean ± S.D. of relative intensity of each Sdk::mScarlet construct at tricellular junctions normalized to intensity at the adjacent bicellular junctions in the pupal dorsal thorax epithelia. *n* = 48 tricellular junctions from four flies each. *p*-value by an unpaired two-tailed Welch’s *t*-test. (E) Representative image of endogenously Venus-tagged Cno (Cno::Venus) in the pupal dorsal thorax epithelia (left) and mean ± S.D. of relative intensity of Cno::Venus at tricellular junctions, quantified similarly to (B). *n* = 36 tricellular junctions from three flies. (F) Representative images of *in vitro* His_6_::ICD::mEGFP condensates in bulk phase (in buffer of 20 mM HEPES pH 7.25, 150 mM KCl) at indicated bulk protein concentrations in the presence (Sdk^ICD^) and absence (Aka^ICD^) of 5% PEG-20,000. Images of Sdk^ICD^ at 12 µM and Aka^ICD^ at 10 µM are identical to those in Figure 1C. (G) Representative image of Sdk^ECD-TMD^-FUS^PLD^::mScarlet with E-Cad::GFP in the pupal dorsal thorax epithelia. Arrowheads indicate some of tricellular junctions.

**Figure S2.**
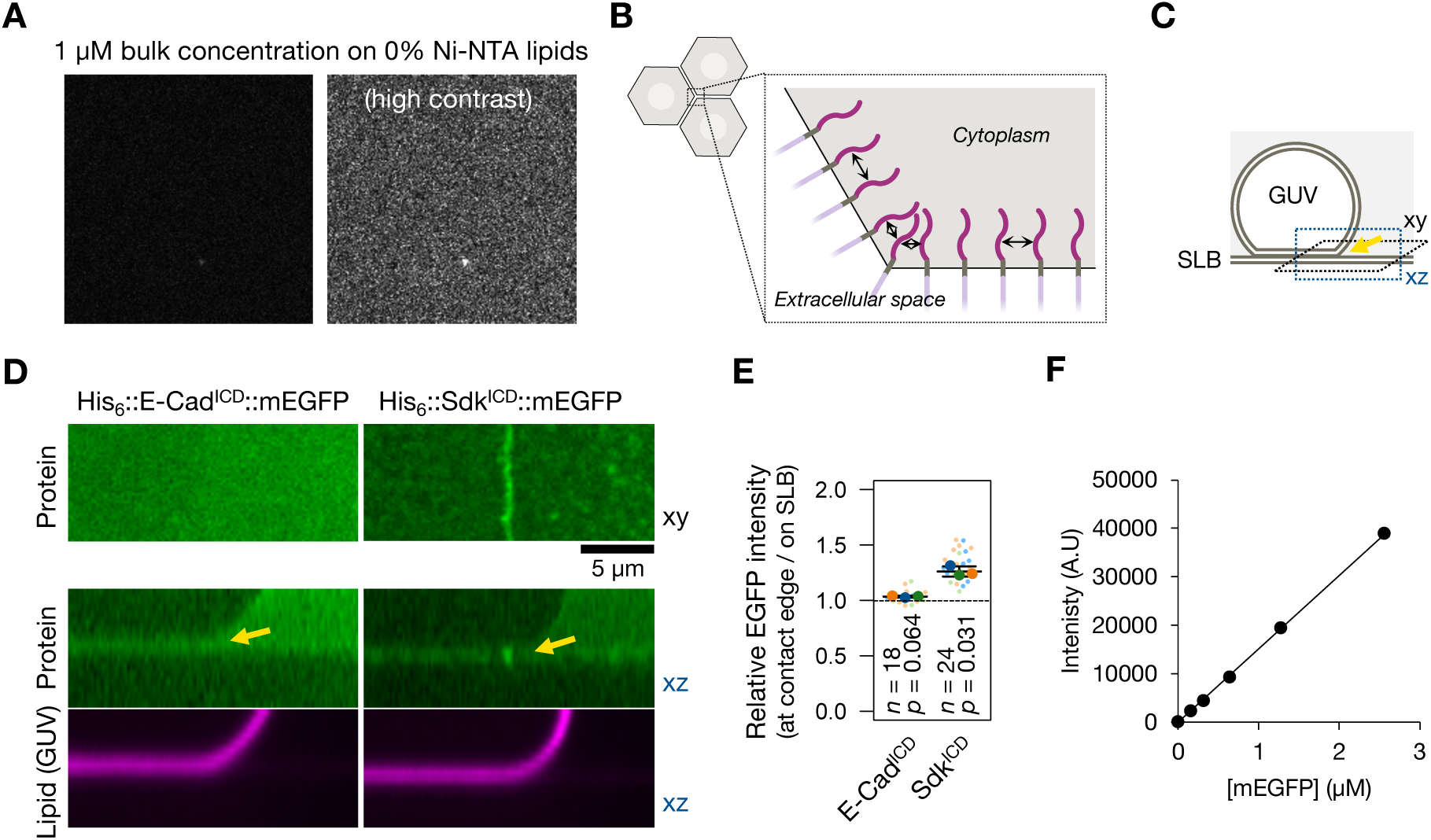
Sdk intracellular domain localizes at angled points of lipid membranes *in vitro*. (A) An identical assay to Figure 2B except the Ni-NTA conjugated lipid concentration (0%). The left image is at the same contrast as in Figure 2B. (B) Simplified schema of a magnified view around tricellular junctions with distribution of single-pass transmembrane proteins on the plasma membrane. Arrows, distance of the adjacent intracellular domains (dark magenta). (C) Schema of *in vitro* protein condensation assay on SLBs and GUVs. Arrows, the edge of an SLB-GUV contact. Proteins are placed outside the GUVs (grey). (D) Representative images of His_6_::ICD::mEGFP (green) at bulk concentration of 1 µM on SLBs and GUVs (magenta) containing 1% Ni-NTA conjugated lipids in buffer (20 mM HEPES pH 7.25, 150 mM KCl, and 1 mM MgCl_2_). Top, planar images at the level of SLBs with one GUV attached; bottom, their cross sections. Note that only lipid signals on GUVs were apparently visible. (E) Mean ± S.D. of relative intensity of His_6_::ICD::mEGFP at the edges of GUV-SLB contacts normalized to that on SLBs in the proximity. *n*, number of the edges examined from three independent experiments. *p*-values by a one-sample *t*-test with a test value of 1. (F) Fluorescent intensity of purified mEGFP proteins imaged in the same microscope setting as in Figure 2C in buffer (20 mM HEPES pH 7.25, 150 mM KCl, and 1 mM MgCl_2_) as a function of the protein concentration. This standard curve was used for estimating EGFP-tagged protein concentrations *in vivo* and *in vitro*.

**Figure S3.**
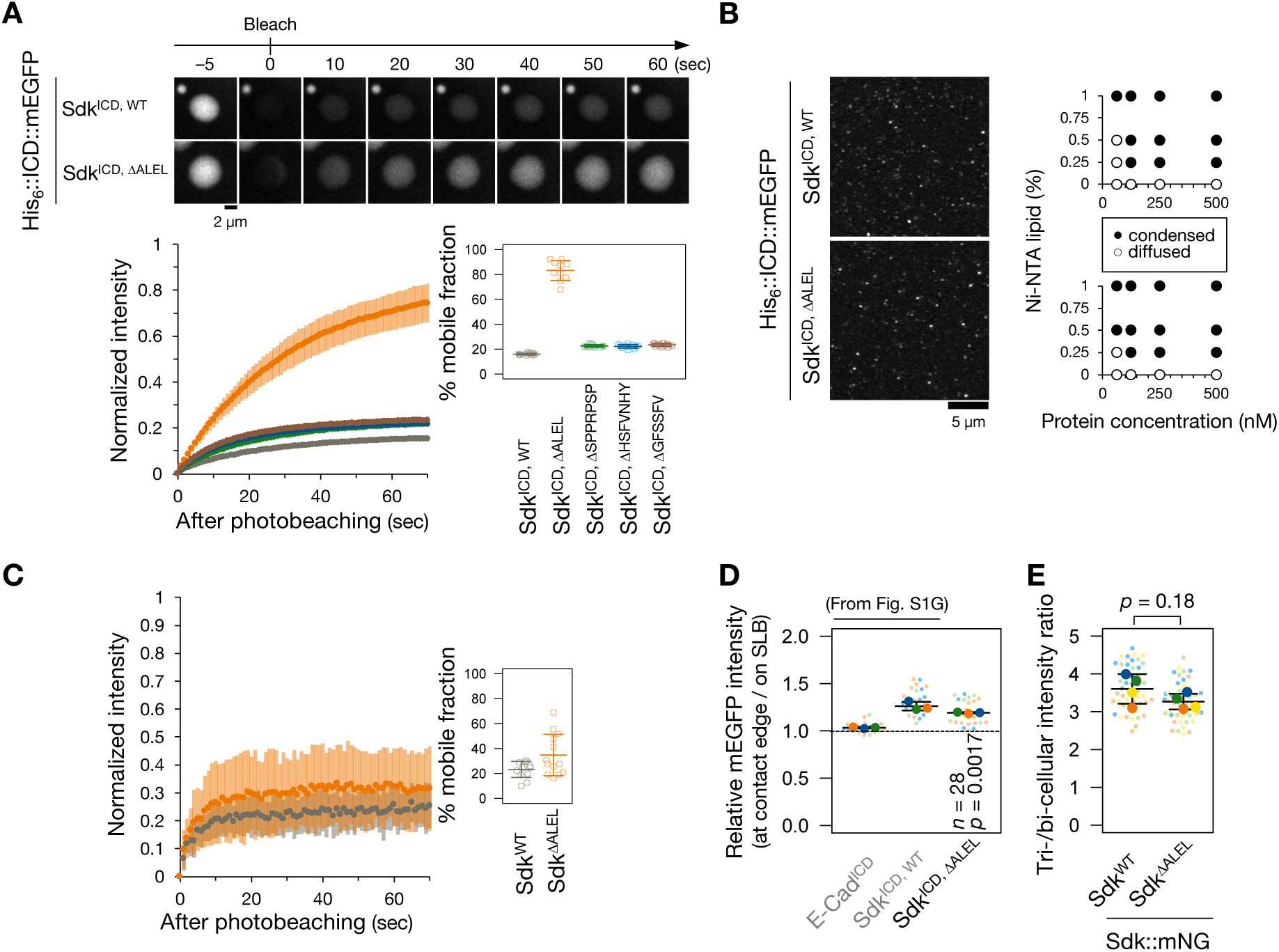
Dynamics of ALEL-deleted Sdk intracellular domain. (A) Top, time-lapse images of one condensate each of His_6_::Sdk^ICD, WT^::mEGFP and His_6_::Sdk^ICD, ΔALEL^::mEGFP in solution before and after photobleaching at the whole condensate areas (0 sec). Bottom, mean ± S.D. of normalized intensity of wild-type and each deletion of His_6_::Sdk^ICD^::mEGFP in condensates after photobleaching (left) and mean ± S.D. of mobile fraction (right). *n* = 10–15 condensates for each construct. Proteins were at 15 µM in buffer (20 mM HEPES pH 7.25, 150 mM KCl, and 5% PEG-20,000). (B) Left, representative images of His_6_::Sdk^ICD, WT^::mEGFP and His_6_::Sdk^ICD,ΔALEL^::mEGFP on SLBs containing 0.5% Ni-NTA-conjugated lipids in buffer (20 mM HEPES pH 7.25, 150 mM KCl, and 1 mM MgCl_2_) at a bulk concentration of 500 nM. Right, 2D charts indicating condensation behaviors of each construct as functions of protein bulk concentrations and Ni-NTA lipid percentages. (C) Mean ± S.D. of normalized intensity of His_6_::Sdk^ICD, WT^::mEGFP and His_6_::Sdk^ICD, ΔALEL^::mEGFP in condensates on SLBs after photobleaching (left) and mean ± S.D. of mobile fraction. *n* = 13 condensates for each construct. (D) Relative intensity of His_6_::Sdk^ICD, ΔALEL^::mEGFP at the edges of contact of GUVs and SLBs normalized to that on SLBs in the proximity. *n*, number of the edges examined from three independent experiments. *p*-values by a one-sample *t*-test with a test value of 1. The other plots are from Figure S2D for comparison. (E) Mean ± S.D. of relative intensity of Sdk^WT^::mNG and Sdk^ΔALEL^::mNG at tricellular junctions normalized to intensity at the adjacent bicellular junctions in the pupal dorsal thorax epithelia. *n* = 48 tricellular junctions from four flies each. *p*-value by an unpaired two-tailed *t*-test.

**Figure S4.**
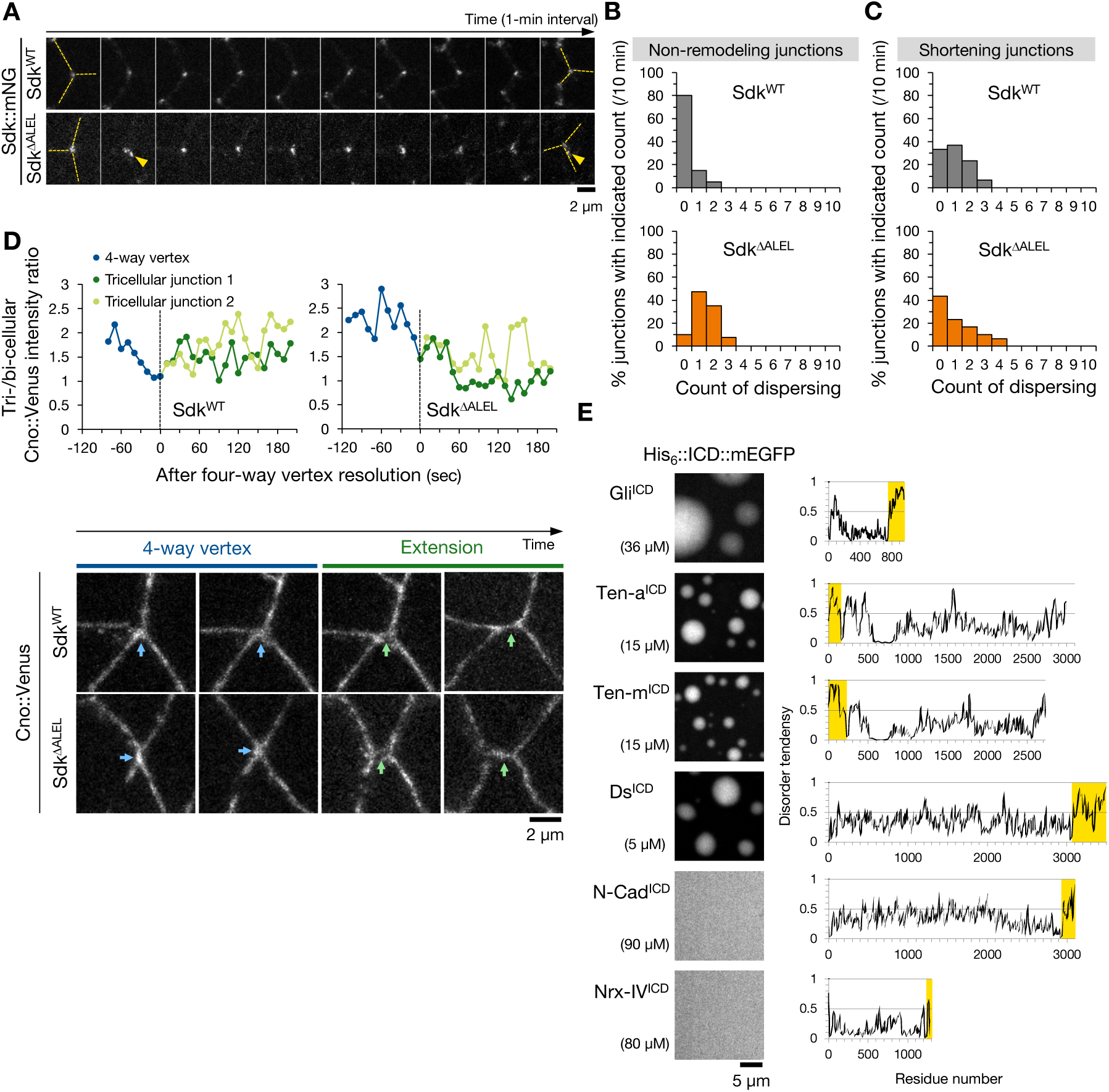
Dynamics of Sdk and Cno in Sdk^ΔALEL^ flies during tissue movement. (A) time-lapse images of Sdk::mNG at tricellular junctions formed by three non-remodeling junctions during germband extension. Broken lines, the three junctions (labeled in the first and last images). Yellow arrowheads, dispersed signals of Sdk::mNG on bicellular junctions. (B) Proportion of frequency of the dispersing events of Sdk::mNG happening at non-remodeling junctions in 10 min. *n* = 40 non-remodeling tricellular junctions from five (WT) and four (ΔALEL) embryos, respectively. *p* = 8.4 × 10^−8^ by two-tailed Mann-Whitney *U*-test. (C) Proportion of frequency of the dispersing events of Sdk::mNG happening on neighboring junctions of shortening junctions in 10 min. *n* = 30 extending junctions from three embryos each genotype. *p* = 0.91 by two-tailed Mann-Whitney *U*-test. (D) Top, relative intensities of Cno::Venus at four-way vertices and the two tricellular junctions (edges) of extending junctions over the neighboring bicellular junctions of each representative remodeling junction in Sdk^WT^::mScarlet and Sdk^ΔALEL^::mScarlet flies. Broken line, the timing of four-way vertex resolution. Bottom, representative images of Cno::Venus during the four-way vertex and junction extension phases. Blue and green arrows, four-way vertices and extending junctions, respectively. (E) Left, representative images of *in vitro* His_6_::ICD::mEGFP or mEGFP::ICD::His_6_ condensates in bulk phase (in buffer of 20 mM HEPES pH 7.25, 150 mM KCl, and 5% PEG-20,000) at indicated bulk protein concentrations each. Intracellular domains were chosen from fly adhesive single-pass transmembrane proteins: Gliotactin (Gli), Tenascin accessory (Ten-a), Tenascin major (Ten-m), Dachsous (Ds), N-Cadherin (N-Cad), and Neurexin IV (Nrx-IV). Right, their disorder prediction on IUPred3. Yellow boxes, the position of each intracellular domain.

Video S1 and S2

Time-lapse images (1-sec interval) of 15 µM of WT (Movie 1) or ΔALEL (Movie 2) His_6_::Sdk^ICD^::EGFP in buffer (20 mM HEPES pH 7.25, 150 mM KCl, and 5% PEG-20,000) after 1 h of inducing their condensation. One condensate each are coming to other condensates on the bottom.

Video S3 and S4

Time-lapse images (10-sec interval) of MRLC::mScarlet-I (magenta) with Sdk^WT^::mNG (Movie 3) or Sdk^ΔALEL^::mNG (Movie 4) (green) around extending cell–cell junctions in germband extension.

